# Structural and dynamic study of fungal cell wall degrading fungal chitinase and its interaction with chitooligosaccharide

**DOI:** 10.1101/2025.06.29.662179

**Authors:** Uttam Kumar Jana, Pratyoosh Shukla, Naveen Kango

**Affiliations:** Department of Microbiology, Dr. Harisingh Gour Vishwavidyalaya (A Central University), Sagar, Madhya Pradesh, India 470003; Enzyme Technology and Protein Bioinformatics Laboratory, School of Biotechnology, Institute of Science, Banaras Hindu University, Varanasi, Uttar Pradesh, 221005, India

**Keywords:** Chitin, chitinase, fungi, molecular simulations, free energy

## Abstract

Chitin, comprising of repeating units of N-acetyl-glucosamine, is the second most abundant polymer occurring in wide range of insects, fungi, yeasts and plants. Chitinases hydrolyze chitin into chitooligomers which finds multifarious uses in various sectors and are gaining attention particularly as a biocontrol agent against chitin-containing insects and plant pathogens. Although fungi are a significant source of chitinases, the phylogenetic and functional diversity of the fungal enzymes is not well understood. In this study, we employed molecular modeling and simulation techniques to investigate the molecular characteristics of chitinase (*PmChi*) from a mycoparasitic fungus, *Paraphaeosphaeria minitans*, widely used as a biological control agent. The secondary structure of *PmChi* is predominantly random coil (51.24%), followed by alpha helices (28.44%) and extended strands (14.22%). *PmChi* contains the conserved chitinase sequence FDGLDIDWE at positions 178 to 186, where glutamic acid (E) acts as the catalytic proton donor, and aspartic acid (D) stabilizes the protein by accommodating substrate distortion. The protein surface of *PmChi* is rich in non-polar amino acid residues, while the active site contains more polar residues to facilitate the reaction. Key amino acids involved in catalytic activity include Trp146, Asp184, Glu186, Tyr187, Pro228, Met252, Tyr254, and Asp255. Molecular simulations demonstrated that *PmChi* maintained stability during interaction with chitotriose. Residual flexibility, hydrogen bonding, and structural packing showed consistent trajectories with no significant perturbations throughout the simulation. The free energy calculations for the *PmChi*-chitotriose complex indicated that MM/PBSA calculations are more accurate for analyzing enzyme-carbohydrate interactions. These findings enhance our understanding of the structural properties and functional dynamics of chitinase from *P. minitans* and provide a platform for future research and applications.

## 1. Introduction

Chitin is a structural biopolymer composed of repeating units of N-acetyl-D-glucosamine linked *via* β-1, 4-glycosidic bond. It is present in insect cuticle as well as cell wall of fungi as a protective structural component to stand with physical abrasion, chemical erosion, and pathogenic invasion (Qu et al., 2021). The molecular composition of chitin is like cellulose, differentiated by the presence of acetamide group (-NHCOCH_3_) present at the C-2 position of glucose unit (Bhagwat et al., 2021). Chitinases (EC 3.2.1.14) are glycoside hydrolases which cleave the β-1, 4-glycosidic bond of chitin biopolymer and liberate mono-sugars and oligo-sugars. Chitinases are classified into five classes belonging to the two glycosyl hydrolase group GH18 and GH19. GH19 are plant based chitinases and found in some *Streptomyces* that reside in class I, II and III. At molecular level, GH18 contain (α/β)_8_ barrel domain for main catalytic activity and GH19 have a bi-lobed configuration dominated by α-helices (Bhagwat et al., 2021). Different subtypes like chitinase-A and B are also present in nature based on the amino acid sequence identity but they have remarkable difference in biochemical and antifungal activities. Likewise, chitinases-A from different sources including fungi been exploited as a biocontrol agent (Liu et al., 2020). The microbial inoculants potentially replacing harmful pesticides are called biocontrol agents. Biological control, by definition, provides a non-chemical method for management of plant diseases by using other living entities, such as microorganisms. The biocontrol capacity of a microbe can result from production of antibiotic compounds, or enzymes capable of fungal cell wall lysis, depletion of iron from the rhizosphere, induced systemic resistance, and competition for niches with pathogens within the rhizosphere. Production of one or more antibiotics is a mechanism most associated with biocontrol ability. Several biocontrol strains can also produce antifungal enzymes, for example chitinases, β1,3-glucanases, proteases, or lipases, with the capacity to lyse fungal cells. Synthesis of low-molecular mass siderophores that chelate iron in the soil near roots can inhibit the proliferation of fungal pathogens. *Paraphaeosphaeria minitans* (formerly known as *Coniothyrium minitans*, a mycoparasite, is a biocontrol agent against *Sclerotinia* spp. Ascomycota, *Sclerotinia* has significant threat to agriculture sector and infect large number of agriculture plant species which can led to serious economic crisis for any country. *Sclerotinia* is responsible for diseases like white mold in over 400 plant species including key crops such as soybeans, sunflowers, canola, and beans (Geffersa et al., 2023). This pathogen can survive in cool as well as in moist environments which results widespread crop losses. Between 1996 and 2009, *Sclerotinia* stem rot was responsible for significant yield losses exceeding 10 million bushels. The most severe losses occurred in 1997, 2004, and 2009, with 35, 60, and 59 million bushels. These losses translated to approximately 227 million USD in 1997, 344 million USD in 2004, and 560 million USD in 2009 (Peltier et al., 2012). Governments may face rising costs for agricultural subsidies and import dependencies. If not managed effectively, fungal outbreaks can destabilize local economies and negatively impact global food supply chains (Springmann & Freund, 2022). The pathogen generates large number of sclerotia on for survival for long time. *P. minitans* could secrete large number of cell wall degrading enzymes encoded by different genes. Chitinase help to invade the *P. minitans* in *Sclerotinia* spp. through parasitical process (Nicot et al., 2019; Wei et al., 2016). Various investigations about fungal chitinases have been reported. Chitinase from *Trichoderma longibrachiatum* was transformed into cotton plants to control *Aphis gossypii*(Anwar et al., 2023). In agriculture, chitinases are being used to naturally control pests and plant fungal diseases (Oyeleye & Normi, 2018). These enzymes have shown success in managing harmful fungi like *Colletotrichum gleosporoides* (de la Fuente Salcido et al., 2016), *Trichoderma reesei* (Taira et al., 2002), *Rhizoctonia solani* (Ghasemi et al., 2010), *Phoma medicaginis*(Slimene et al., 2015), *Fusarium graminarium*(Hjort et al., 2014) and *Fusarium oxysporum* (Bhattacharya et al., 2016).

There is limited information available in the literature against the structural and functional properties of *P. minitans* chitinases. Native three-dimensional structure is needed to understand the biological and functional properties of chitinase. X-ray crystallography is a popular technique for analyzing the protein structure but remains limited due to high cost and technical expertise requirements (Davis et al., 2008). In recent years, *in-silico* study has achieved significant prediction of protein molecules to understand different biological processes. Computational techniques give insight of protein folding and determine catalytic sites of the protein. In present report, homology modeling, molecular docking, and molecular dynamics simulations of chitinase from *P. minitans* were performed for analyzing the physical parameters, domain, and motif and three level structural analysis. Interaction with chitooligosaccharide is also reported which will help to gain knowledge about chitinase from mycoparasite for future protein engineering in agriculture applications.

## 2. Materials and methods

### 2.1 Sequence and phylogenetic analysis

Protein sequence information of chitinase of *P. minitans* was collected from UniProt database (http://www.uniprot.org/) with Uniprot ID (Q9HGU5). Conserved domains present in two enzymes were analyzed using the Pfam (Mistry et al., 2021), InterProScan (Quevillon et al., 2005) and <u>C</u>onserved <u>D</u>omain <u>D</u>atabase (CDD) database (J. Wang et al., 2023). Protein phylogenetic tree was determined using the Blast Tree View of ten highest identity protein from BlastP results. The evolutionary history of chitinase from *P. minitans* was analyzed by fast minimum evolution tree method. Evolutionary distance between different species chitinases was evaluated using expected fraction of amino acid substitutions per site, given the fraction of mismatched amino acids in the aligned region (Grishin, 1995).

### 2.2 System preparation, refinement, and validation

Homology modeling of chitinase from *P. minitans* (*PmChi*) was performed using the RaptorX (S. Wang et al., 2016). Homology models were retrieved based on the highest alignment score and distance-based protein folding powered by deep learning. Hydrogen bonding and the energy minimization of the modeled protein was optimized using the DeepRefiner (Shuvo et al., 2021). SOPMA was used for the analyzing the protein secondary structures (Geourjon & Deléage, 1995). Procheck was employed for the analyzing stereo-chemical quality of the model residue-by-residue using Ramachandran plot (Laskowski et al., 1993). The comparison between X-ray diffraction and NMR structure was predicted by the ProSAweb (Wiederstein & Sippl, 2007a). Statistical analysis of different atoms present in the model and their non-bonded interaction was analyzed using the ERRAT (Colovos & Yeates, 1993). Verify 3D was applied for the calculation of compatibility between atomic model and the amino acid sequence by estimating the model under different location and environment (polar, non-polar, loop, alpha, beta, etc)(Eisenberg et al., 1997a). The secondary and super secondary model of chitinase from *P. minitans* was calculated by Stride. Different other parameters like interpolated charges, hydrophobicity, hydrogen bonds, aromatic surface, solvent accessibility, and ionizability of chitinase was evaluated using Discovery Studio Visualizer. The model is available in ModelArchive at https://www.modelarchive.org/doi/10.5452/ma-lnvv8.

### 2.3 Theoretical characteristics of *PmChi*

Different physicochemical characters such as molecular weight (Mw), number of positive and negatively charged amino acids, extinction coefficient, theoretical isoelectric point (pI), aliphatic index, instability index and grand average hydropathicity (GRAVY) of *PmChi* was predicted by ExPASy’s ProtParam web server tool (http://web.expasy.org/protparam/). Secondary structure, solvent accessibility, and different ontology studies, such as molecular function, cellular components, and biological process were predicted by Predict Protein server (https://predictprotein.org/).

### 2.4 Ligand preparation

Chitotriose canonical SMILES sequence was retrieved from the PubChem (CID 121978) and 3D structure was constructed using the UCSF chimera(Pettersen et al., 2004). Energy of the chitotriose was minimized using the minimize structure tools, where the parameters like steepest descent steps (100), steepest descent step size (0.02 Å), conjugate gradient steps (10), conjugate gradient step size (0.02 Å) and update interval (10) were set for the minimization step. The minimizations of standard residues were performed using the AMBER ff14SB force field and other residues were executed using the Gasteiger force field.

### 2.5 Molecular docking

The interaction *PmChi* with chitotriose was analyzed using the molecular docking method. The molecular docking was performed using the AutoDock 4.2 (Morris et al., 2009). The grid box was generated with the help of AutoGrid and selected 3D coordinates in x, y, z-dimensions, where the active site of the enzyme as well as a large portion of adjoining surface was covered. After successful run, possible chitotriose docking structure containing the protein-ligand complex were generated and best complex was selected based on minimum binding energy (Kcal/mol), hydrogen bonds numbers, docking score, and weak interactions for the further study. Results were visualized using Discovery Studio Software. Absolute binding affinity of *PmChi*-chitotriose complex was predicted by K_DEEP_ (Jiménez et al., 2018).

### 2.6 Molecular Dynamics (MD) Simulation

MD simulations of modelled *PmChi* and *PmChi*-chitotriose complex were conducted with Gromacs 2020 (http://www.gromacs.org/) using CHARMM27 force field I periodic boundary conditions on the local GPU server. The structures were solvated in a truncated octahedron box of simple point charge water model. The solvated system was neutralized with Na^+^ or Cl^−^ counter ions using the tleap program. Particle mesh Ewald was employed to calculate the long-range electrostatic interactions. The cut-off distance for the long-range Van der Waals energy term was 12.0 Å. The system was then minimized at a maximum force of 1000.0 KJ/mol/nm using 50,000 steps. The solvated and energy minimized systems were further equilibrated for 100 ps under NVT and NPT ensemble processes. The dynamics were integrated using the velocity Verlet integrator, with a time step of 2 fs and bonds constrained using the LINCS algorithm. The simulations were performed for 200 ns and PDB structure of the complex were retrieved at every 25 ns. The results were analyzed using Gromacs toolkits gmx rms, gmx rmsf, and gmx gyrate tools for RMSD, RMSF, and Rg respectively.

### 2.7 Determination of Binding Free Energies

To determine the binding free energy, we utilized the gmx_MMPBSA, which combines GROMACS and AmberTools functions for performing end-state free energy calculations(Valdés-Tresanco et al., 2021). This tool calculates the average molecular mechanical energy, considering both bonded (bond, angle, dihedral) and nonbonded interactions (Van der Waals and electrostatic interactions), using molecular mechanics (MM) force-field parameters. This approach allows us to break down the binding energy into contributions from each residue, providing a more comprehensive understanding of the binding mechanisms. In our study, we performed a MM/GBSA analysis on the trajectories obtained from molecular dynamics simulations of *PmChi*-chitotriose complex with a focus on the contributions of all amino acids located within 6 Å from the binding interface to the binding energies. Energy calculations were performed using the MMPBSA.py core. Results were analyzed and visualized with gmx_MMPBSA_ana tool.

### 2.8 Intra-atomic interaction of *PmChi*

The non-covalent intra-molecular interaction including H-bonds, ionic bonds, disulfide bonds, π-cation, and π-π stacking bonds of the *PmChi* at the atomic level was visualized using the Residue Interaction Network Generator (RING) web server and Arpeggio web server(Jubb et al., 2017; Piovesan et al., 2016). The salt bridges in the protein structure were analyzed by ESBRI web server (Kumar & Nussinov, 1999).

## 3. Results and discussions

### 3.1 Sequence and phylogenetic analysis

Chitinase from *P. minitans* clustered by *P. sporulosa* chitinase with 95.71% similar identity and its conserved domain is shown in the Figure 1A. Phylogenetic analysis showed the similarity of chitinase sequence with other proteins were to be 93.36% (*Didymosphaeria variabile*, XP_056074141.1), 93.00% (*Karstenula rhodostoma*, KAF2438673.1), 84.27% (*Binuria novae zelandiae*, KAF1975465.1), 79.80% (*Lophiostoma macrostomun*, KAF2656671.1) and 74.04% (*Lentithecium fluviatile*), respectively. All chitinases belonged to GH18 family which indicated fungi possess many GH18 coded genes and chitinase had chitin binding domains evolved through gene expansion and contraction many times during this protein evolution (Hartl et al., 2012).

**Figure 1:**
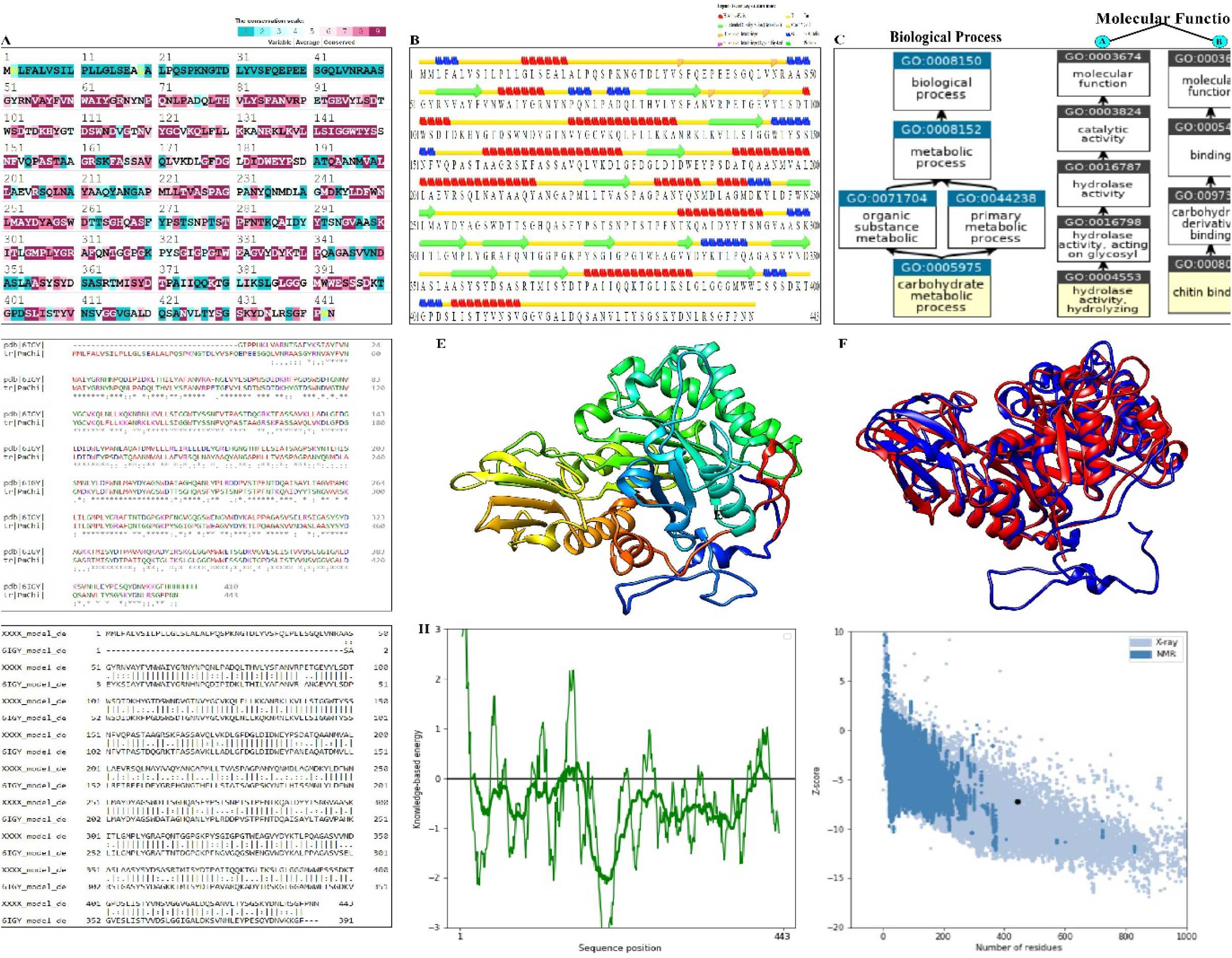
Model generated for *P. minitans* chitinase (*PmChi*) and its analysis. (A) Conservation at the level of residue of *PmChi* denoting that the enzyme belongs to GH18 carbohydrate active enzyme family, (B) Predicted secondary structure of *PmChi* show majorly α-helices followed by extended strand and β-turn were the main structure components as revealed by Stride, (C) Gene ontology analysis for *PmChi* in case of molecular function and biological process, (D) Basepair wise conserved domain analysis of *PmChi* with structure alignment of *PmChi* and chain A of chitinase from *Aspergillus niger* (PDB ID: 6IGY), (E) The Deeprefiner energy minimized model of *PmChi* by UCSF Chimera, (F) Image of superpose structure between *PmChi* and chain A of 6IGY, (G) Basepair wise structure alignment of *PmChi* and chain A of 6IGY, (H) Protein Structure Analysis of *PmChi*. Overall model quality and local model quality of the enzyme model showing most of the sequence in the negative energy mode.

### 3.2 Theoretical characteristics of *PmChi*

The physico-chemical properties of chitinase were computed by ProtParam tool and found that the molecular weight of *PmChi* was around 47 kDa (Table 1). Theoretical pI of the chitinase was 4.96 which denoted that overall charge of the protein was negative because of the relative amount of acidic amino acid present in *PmChi*. Similarly, IFF electrophoresis of a chitinase from black soybean seeds showed isoelectric point at 4.34 (Murakami et al., 2016). *PmChi* contained high number of acidic (negatively charged) amino acid residues than basic amino acids (positively charged). Previously, it was also reported that chitinases from *Beauveria bassiana* had the similar richness of acidic amino acid in its backbone. Large number of charged residues in protein have ability for the variation of activity and it could be potential mutant sites for changing the enzymatic function (Wu et al., 2020). *PmChi* had high protein solubility for efficient degradation of the polymer as aspartic acid and glutamic acid contribute favorably to protein solubility compared to other amino acids (Trevino et al., 2007). Instability index is used to measure the half-life of a protein *in vivo* condition. Instability index of *PmChi* based on sequence-specific elements was 32.60 which suggested that stability of the protein *in vivo* condition is stable as the value was lower than 40. Previously, it was established that unstable proteins had > 40 value whereas all the stable proteins had < 40 value (Guruprasad et al., 1990). The predicted GRAVY value of *PmChi* was negative as shown in the Table 1. It means that *PmChi* tends to be more hydrophobic which supported our previous conclusion as protein has more acidic amino acids and would be more charged. The aliphatic index of *PmChi* was 74.94 that shows *PmChi* have the thermostability, as increase in aliphatic index refers to thermostability of globular protein (Ikai, 1980).

**Table 1.** Predicted physico-chemical characteristics of chitinase (*PmChi*) from.

|  |  |
| --- | --- |
| Protein ID (Uniprot) | Q2TXJ2 |
| Sequence length (No. of AA) | 443 |
| Molecular weight | ~47 KDa |
| pI | 4.96 |
| Total number of negatively charged residues<br>(ASP + GLU) | 36 |
| Total number of positively charged residues<br>(ARG + LYS) | 28 |
| Extinction coefficient | 99700 |
| Instability index (II) | 32.60 |
| Aliphatic index | 74.94 |
| GRAVY | -0.249 |

*Paraphaeosphaeria minitans* through ProtParam tool

### 3.3 Secondary structure analysis

The secondary structure of *PmChi* is dominated by random coil region (51.24%) followed by alpha helix (28.44%) and extended strand (14.22%) (Figure 1B). The secondary structure rich in random coils and alpha-helix regions gives stability to the protein and random coil present in between alpha helix and beta strand tend to be less as residues compared to other types of random coils(Khrustalev, 2020). In case of post-translational modification of *PmChi*, there is two Asparagine-N glycosylation sites that were present (Asn-Xaa-Ser/Thr sequons) at 27 and 276 positions. Glycosylation helped *PmChi* to get secreted in the extracellular environment. In addition, it also supported other parameters like proper folding, oligomerization, solubility, and protein stability (dos Reis Almeida et al., 2011). Protein signature recognition methods resolved that *PmChi* belonged to glycosyl hydrolase family 18 (GH18) from amino acid range 52-416. Further, subfamily analysis denoted that it was chitinase II, mainly ranging from amino acid region 52 to 398. Group II chitinase is a widespread chitinolytic enzyme, containing greatest number of catalytic domains, have widespread activities such as chitinase, chitodextrinase and killer toxin of *Kluyveromyces lactis* (Chen et al., 2018; Colussi et al., 2005). Homologous superfamily analysis showed that it has chitinase insertion domain (CID) from 307 to 369. Mainly, CID present in chitinase is composed of five or six anti-parallel β-strands and one α-helix insert between TIM barrel (Li & Greene, 2010). Catalytic domain of *PmChi* has (β/α)_8_ TIM-barrel structure where eight internal parallel β-strands are connected by eight α-helices. The active site motif sequence DxxDxDxE for GH18 is important for performing the catalytic reaction and *PmChi* has F<u>D</u>GL<u>D</u>I<u>D</u>W<u>E</u> at the 178 to 186 positions, where E (Glu) is acting as catalytic proton donor and D (Asp) responsible for the stabilization of protein caused by substrate distortion (Tsuji et al., 2010). According to the gene ontology study, *PmChi* belongs to biological process GO:0005975 (carbohydrate metabolic process), have two molecular functions, GO:0004553 (hydrolyzing O-glycosyl compounds) and GO:0008061 (chitin binding) (Figure 1C).

### 3.4 Tertiary structure analysis

Three-dimensional structure of *PmChi* had maximum homology with the chain A of chitinase from *Aspergillus niger* (PDB ID: 6IGY) having 63.43% of identity (Figure 1D). After protein blast with the PDB database, different score such as total scores, query coverage, and E-value between two sequences were 536, 97 % and 0, respectively. These results confirmed the model is suitable for further study. The prepared model was refined by the Deeprefiner and the refined *PmChi* is shown in the Figure 1E. Superpose analysis revealed that RMSD value between two template was 2.77 Å in which 391 atoms in alpha carbon of templates *PmChi* and 6IGY were superposed (Figure 1F). The high RMSD value of superimposition is due to structural difference in the N-terminal region of both proteins. The reason may be species specific differences in chitinase. In Figure 1G, base pair wise two template superimposition is also given. The validation of *PmChi* is also done by ProSA where main parameter Z-score denote the overall structure quality and validation the experimental model for further study. Z-score of *PmChi* structure is -7.21 that denotes the maximum amino acid sequence are negative energy mode (Figure 1H). The negative nature supports the correctness of the model as positive value of the sequence led to problematic and erroneous model(Wiederstein & Sippl, 2007b). QMEAN score was found to be 0.65 ± 0.05, compared with a non-redundant set of PDB structure and found suitable for the study (Figure 2A). 3D protein model was used to perform the compatibility test of the model with its own amino acid sequence as measured by a 3D profile and it was performed by Verify 3D (Eisenberg et al., 1997b). *PmChi* structure was also verified by Verify 3D and it was found that 91.65% of the residues had 3D - 1D score >= 0.2 in *PmChi* after model assessment (Figure 2B). The verification of non-bonded interactions between different atom types was calculated by ERRAT. The ERRAT value for *PmChi* is found to be 90.824. In Figure 2C, there are two error axes with confidence level of 95% and 99%. The Ramachandran plot for *PmChi* revealed that 79.6% of residues were in the most favoured regions, 7.5% in additional allowed regions and 2.1% in generously allowed regions. Ramachandran plot signifies the suitability of the model with respect to the high-resolution structure references (Sobolev et al., 2020).

**Figure 2:**
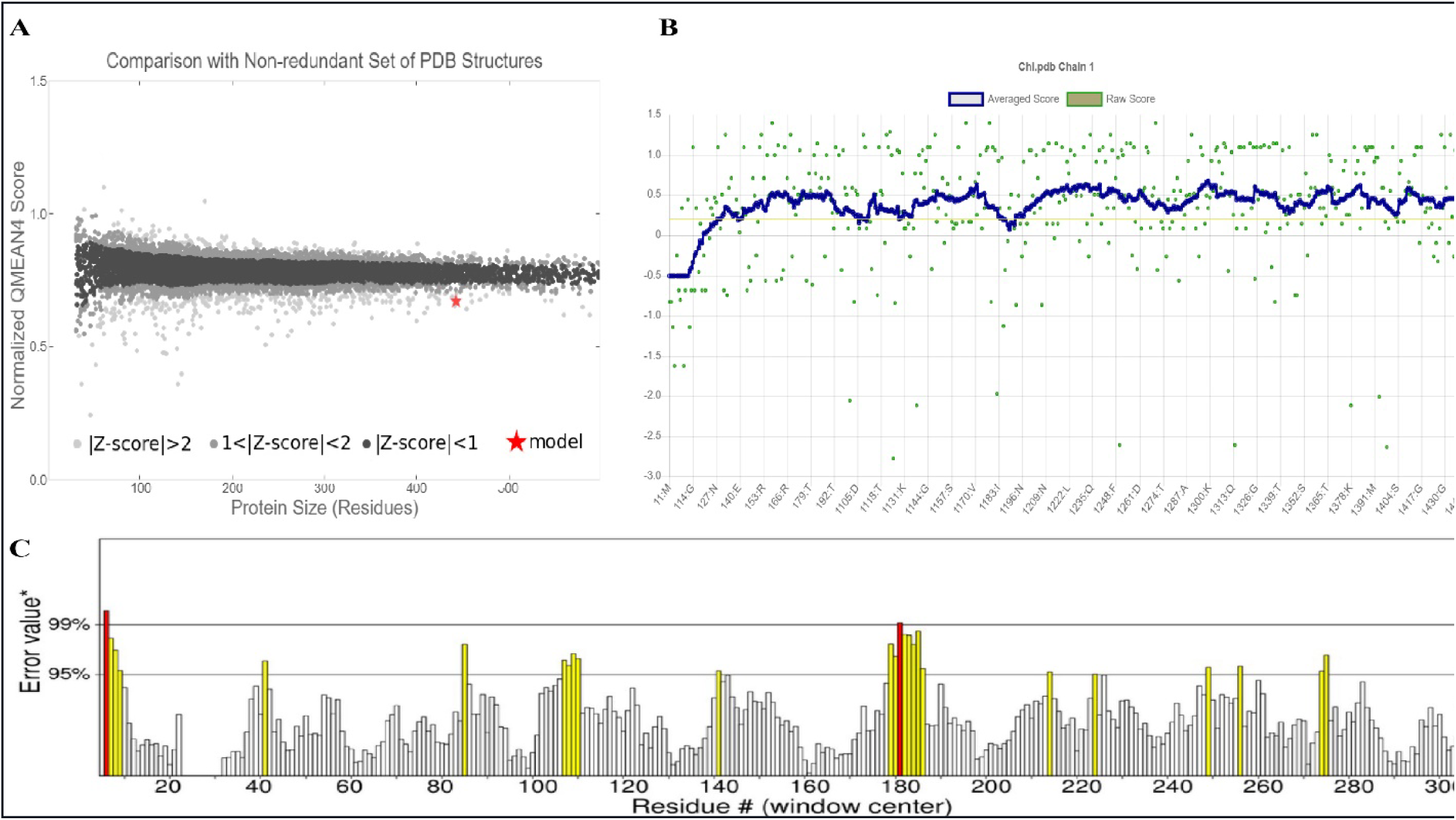
Validation of the predicted *PmChi* model. (A) QMEAN score of the predicted model and local quality estimation of the model, (B) Assessment of 3D model of *PmChi* by Verify 3D and (D) Assessment of non-bonding interaction of different atoms by ERRAT

### 3.5 Docking analysis

The interaction between the chitotriose and the enzyme *PmChi* was investigated using by AutoDock 4.2, and visualized with Discovery Studio. The docking studies identified key amino acid residues involved in the catalytic activity of *PmChi*, specifically Trp146, Asp184, Glu186, Tyr187, Pro228, Met252, Tyr254, Asp255, Trp393, Arg310, and Glu395. These residues are critical for the enzyme’s function, interacting with chitotriose through a variety of chemical bonds, which suggested that a well-defined active site is necessary for substrate binding and catalysis. The docking analysis provided significant insights into the binding affinity and stability of the *PmChi*-chitotriose complex. The Gibbs free energy (Δ*G*) for binding was calculated to be -6.62 kcal/mol, indicating a spontaneous binding process and a stable complex formation. This negative Δ*G* value reflects a favourable interaction between the enzyme and the substrate, which is essential for efficient catalytic activity. Furthermore, the ligand binding efficiency was found to be -0.196 kcal/mol. This value suggests a moderate binding efficiency, implying that while the interaction is stable, there might be potential for optimization to enhance binding affinity. The pKd value of 4.94 supports this, indicating a moderate dissociation constant, which aligns with the observed binding efficiency. The pIC50 value for the docked complex was determined to be 5.169. The pIC50 value is an important parameter in enzyme kinetics and inhibitor studies, as it reflects the potency of an inhibitor. In this context, the pIC50 value suggests a moderate inhibitory potential, providing a basis for further refinement of substrate or inhibitor design to achieve better binding characteristics and catalytic efficiency. Overall, these findings contribute to a deeper understanding of the molecular interactions between chitotriose and *PmChi*, offering a foundation for future studies aimed at improving enzyme-substrate affinity and catalytic performance through rational design and engineering.

### 3.6 Molecular dynamics simulation

#### 3.6.1 Root Mean Square Deviation (RMSD) analysis

The primary objective of the molecular dynamics (MD) simulations was to evaluate the stability of the protein-ligand complex post-docking. To achieve this, the root-mean-square deviation (RMSD) was calculated over the course of the 200 ns simulation. The RMSD value for *PmChi* offers insights into its structural conformation throughout the MD run, while the complex’s RMSD value provides an understanding of its stability during the protein catalytic activity. RMSD value for *PmChi* stabilized at ∼1 nm, whereas that of *PmChi*-chitotriose was found to be ∼0.5 nm (Figure 3A). In the initial phase of the simulation, the *PmChi* protein exhibited fluctuations in its RMSD values from 0 ns to 80 ns. This indicates changes in its structural conformation during this period. However, these fluctuations were absent in the *PmChi*-chitotriose complex, suggesting a more stable structure. The enhanced stability of the *PmChi*-chitotriose complex can be attributed to the binding interactions between chitotriose and the active site amino acid residues of the protein. When chitotriose binds to these residues, it likely forms stable interactions such as hydrogen bonds, hydrophobic interactions, and possibly Van der Waals forces. These interactions can stabilize the overall conformation of the protein, reducing its structural fluctuations. This stabilization effect underscores the significance of ligand binding in maintaining the structural integrity of proteins during catalytic processes. The stability of the protein-ligand complex is crucial for the proper functioning of the protein, as it ensures the active site remains appropriately structured to facilitate catalysis. Thus, the absence of RMSD fluctuations in the *PmChi*-chitotriose complex highlights the potential role of ligand binding in stabilizing the active conformation of the *PmChi*.

**Figure 3:**
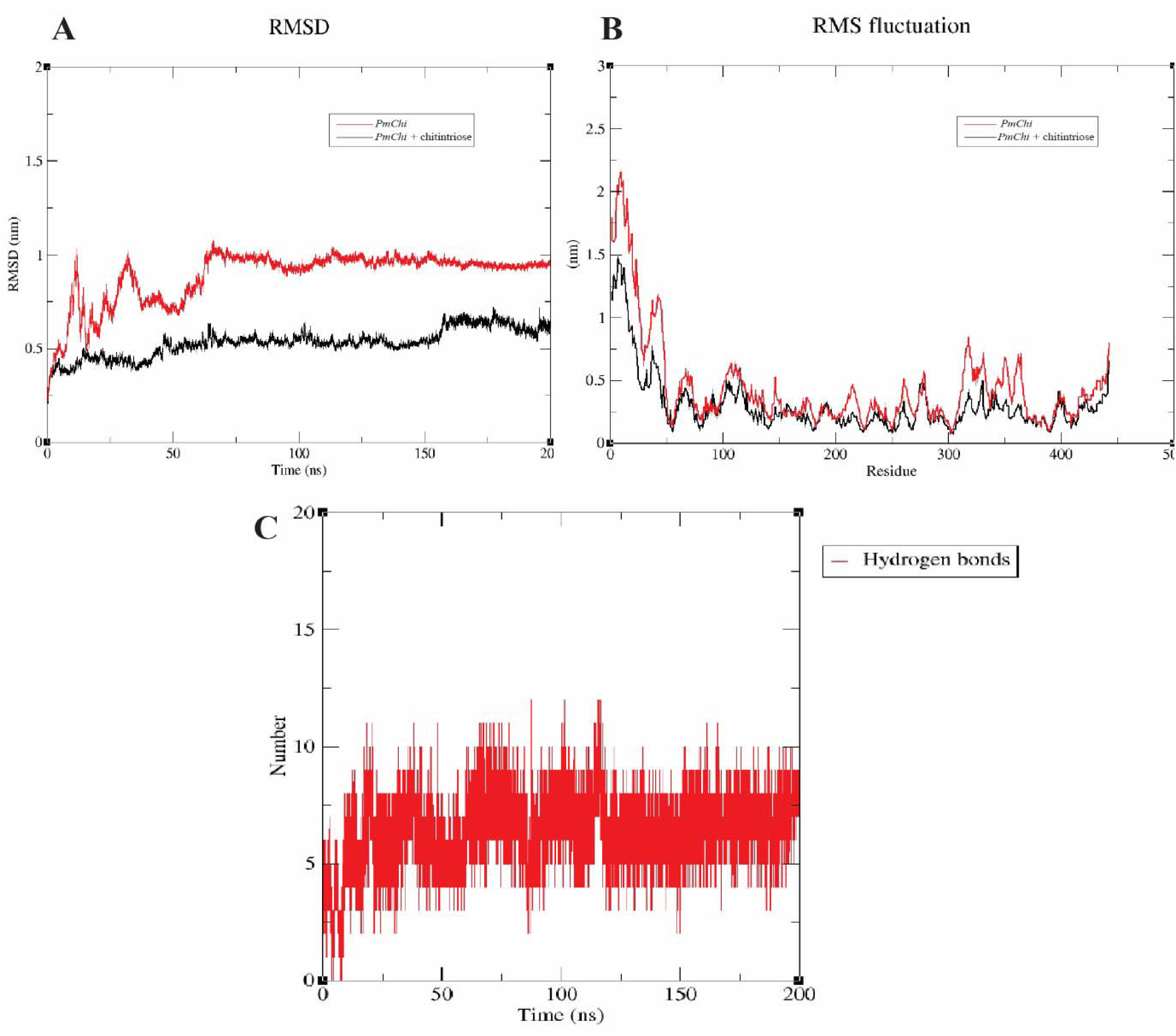
Analysis of different protein dynamics parameters. (A) Time dependent backbone RMSD values with respect to the starting structure of protein and protein ligand complex during MD simulations, (B) Combined RMSF plot and (C) Number of total H-bonds interacting with chitotriose during simulation. Color illustration: Native Protein designated as *PmChi* (red) and *PmChi*-chitotriose complex (black).

#### 3.6.2 Residual flexibility and hydrogen bond analysis

Root Mean Square Fluctuation (RMSF) analysis was conducted to examine the dynamic behavior of the protein structure throughout the simulation trajectory. RMSF values were calculated for each residue in the protein (Jana et al., 2022). Figure 3B displays the RMSF profiles for both *PmChi* and the *PmChi-*chitotriose complex across specific residues. The x-axis indicates the residue number, and the y-axis represents the RMSF values in nanometers (nm). These values were calculated by measuring each residue’s positional deviation from its average position during the simulation. In the case of *PmChi*, all residues exhibited higher RMSF values, indicating increased flexibility compared to other regions. This increased flexibility is crucial for the protein’s function, as it can facilitate the conformational changes necessary for binding interactions or enzymatic activity. Conversely, the *PmChi-*chitotriose complex showed relatively low RMSF values, suggesting a more rigid conformation. This rigidity likely results from the stabilization effect of chitotriose binding, which can restrict the protein’s dynamic movements, leading to a more stable structure(Hashmi et al., 2024). The contrast in RMSF values between the free protein and the complex highlights the impact of ligand binding on protein dynamics, emphasizing the importance of structural flexibility and rigidity in protein function and interaction. Hydrogen bonding calculations are crucial assessments for determining enzyme activity, providing essential insights into the potency and binding strength of interacting molecules. By analyzing hydrogen bonds, researchers can elucidate the stability and specificity of enzyme-substrate or enzyme-product interactions, which are vital for catalytic efficiency. This method has been extensively applied to understand enzymatic mechanisms, offering detailed perspectives on how substrates bind to active sites, undergo catalysis, and transform into products. H-bond was analyzed for the *PmChi-*chitotriose complex using a standard GROMACS hbond tool as shown in Figure 3C. The enzyme carbohydrate interaction was stabilized with chitotriose with four to ten H-bonds in the 200 ns run. Tyr254 and Trp185 are the major amino acids that interact chitotriose with H-bond.

#### 3.6.3 Solvent accessibility

The solvent-accessible surface (SAS) of a protein provides insights into the spatial arrangement of its residues, distinguishing between those exposed on the surface and those buried within the hydrophobic core. This measure is crucial for understanding protein stability, folding, and interactions with other molecules. In the study presented, Figures 4A to 4E depict the time-dependent changes in SAS, specifically in relation to the interaction with chitotriose within the enzyme cavity. Initially, at 0 ns, the enzyme cavity exhibits a high SAS, indicating that many residues are exposed to the solvent (Figure 4A). This high exposure suggests that the enzyme’s active site or binding pocket is accessible, which is essential for substrate binding and catalytic activity. Over time, the SAS within the enzyme cavity decreases. This reduction implies that residues initially exposed to the solvent become less accessible, possibly due to conformational changes in the protein structure. These changes could result from the binding of chitotriose, causing the protein to adopt a more closed or compact conformation. Such a transition often enhances the stability of the enzyme-substrate complex and can be crucial for the proper function of the enzyme. The decrease in SAS also indicates the potential burial of hydrophobic residues that were initially on the surface. This burial can contribute to the stabilization of the enzyme structure and its interaction with the substrate. Understanding the dynamics is vital for elucidating the mechanisms of enzyme action and can inform the design of inhibitors or modified enzymes with improved stability and activity.

**Figure 4:**
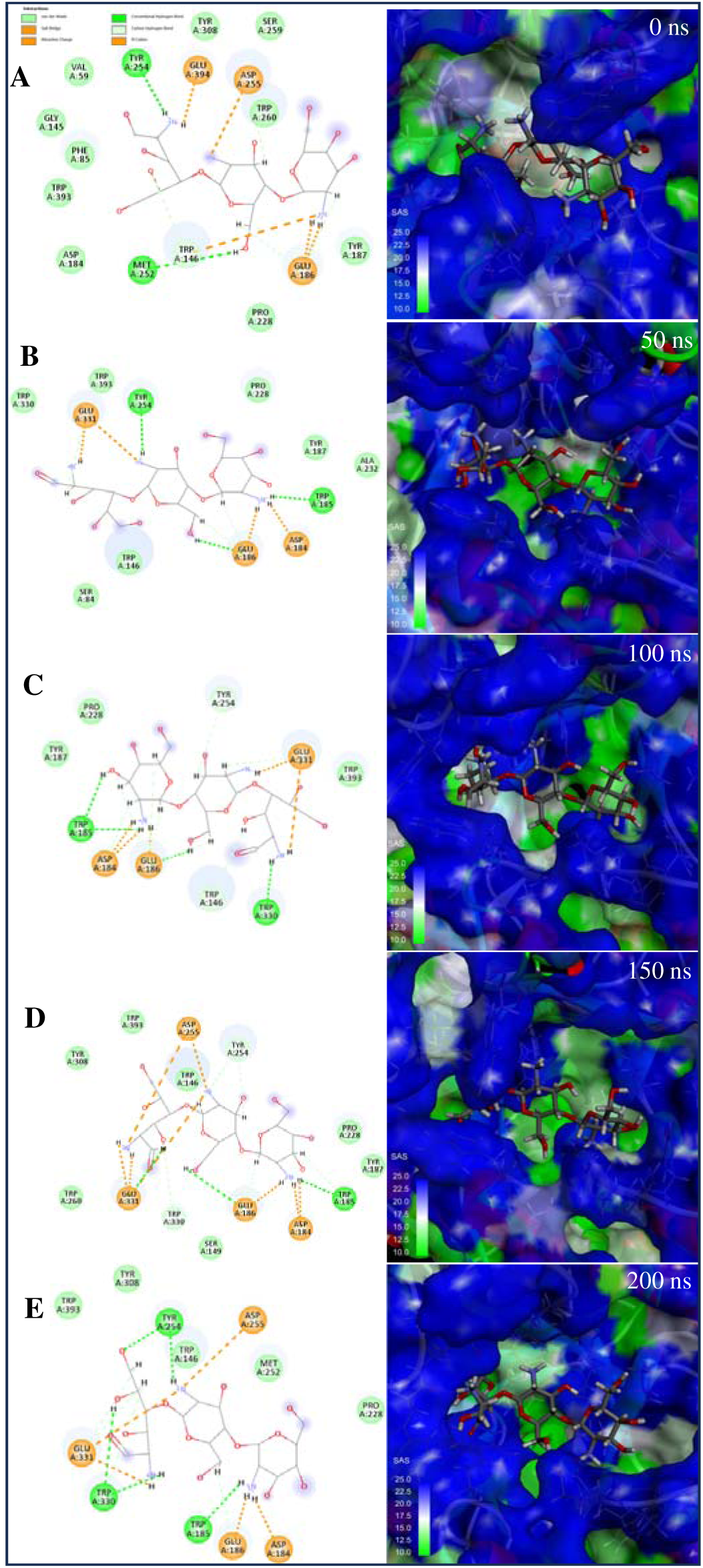
Analysis of solvent-accessible surface during protein dynamics. Time-dependent changes in SAS, specifically in relation to the interaction with chitotriose within the enzyme cavity (A) 0 ns, (B) 50 ns, (C) 100 ns, (D) 150 ns, and (E) 200 ns.

#### 3.6.4 MM/PB(GB)SA analysis

Computational methods integrating molecular mechanics energy with implicit solvation models, like Molecular Mechanics/Poisson-Boltzmann Surface Area (MM/PBSA) and Molecular Mechanics/Generalized Born Surface Area (MM/GBSA), have been extensively used in free energy calculations(Hou et al., 2011). The MM/GBSA and MM/PBSA calculations were carried out for the *PmChi*-chitotriose complex during the molecular dynamic simulations. Figure 6 shows the interacting amino acids and the free energy distribution. In case of MM/GBSA, individual energy contribution was calculated which shows the residues Trp146, Asp184, Glu186, Tyr187, Pro228, Met252, Tyr254, Asp255 and Trp393. The total energy contribution of these residues showed free energy of approximately -4.0, -2.0, -7.0, -0.5, -1.0, -0.8, -3.5, -4.0, and -1.0 kcal/mol respectively (Figure 5A). The framewise energy decomposition graphs for the GB and PB models were similar, suggesting that both methods yield comparable insights into the energetic contributions of individual frames within a molecular system (Figure 5B & D). In case of MM/PBSA, individual energy contribution was calculated which shows the residues Trp146, Asp184, Glu186, Tyr187, Pro228, Met252, Tyr254, Asp255, Trp393, Arg310 and Glu395. The total energy contribution of these residues showed free energy of approximately -2.0, -0.5, -2.7, - 0.2, -0.1, -0.25, -1.25, -2.7, -0.2, +1.25 and -1.9 kcal/mol respectively (Figure 5C). The PB model is theoretically more rigorous than the GB models, making MM/PBSA generally considered superior to MM/GBSA for predicting binding free energies(Hou et al., 2011). All residues exhibited favorable binding energy, except for Arg310, which bound unfavorably with chitotriose. (Figure 5C). Overall, the MM/PB(GB)SA analysis offered valuable insights into the binding free energy and key molecular interactions between the protein and ligand. These findings improve our understanding of enzyme-oligosaccharide interactions and pave the way of future enzyme engineering efforts.

**Figure 5:**
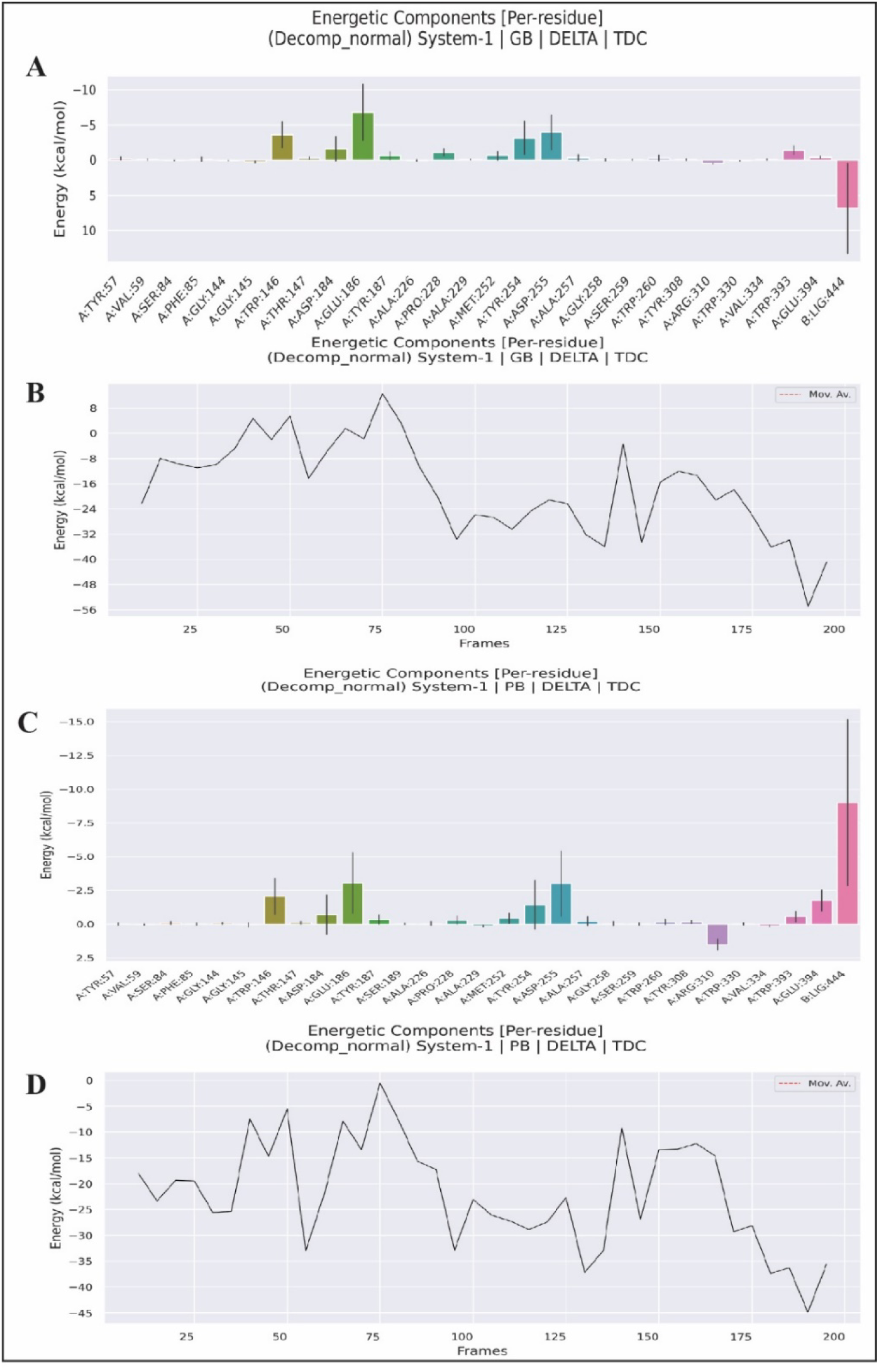
Binding free energy analysis of *PmChi*-chitotriose complex. (A) Generalized Born Surface Area analysis per residues involve in interacting with chitotriose, (B) Frame wise energy graph of *PmChi*-chitotriose complex in Generalized Born Surface Area analysis, (C) Poisson-Boltzmann Surface Area analysis per residues involve in interacting with chitotriose, (D) Frame wise energy graph of *PmChi*-chitotriose complex in Poisson-Boltzmann Surface Area analysis.

**Figure 6:**
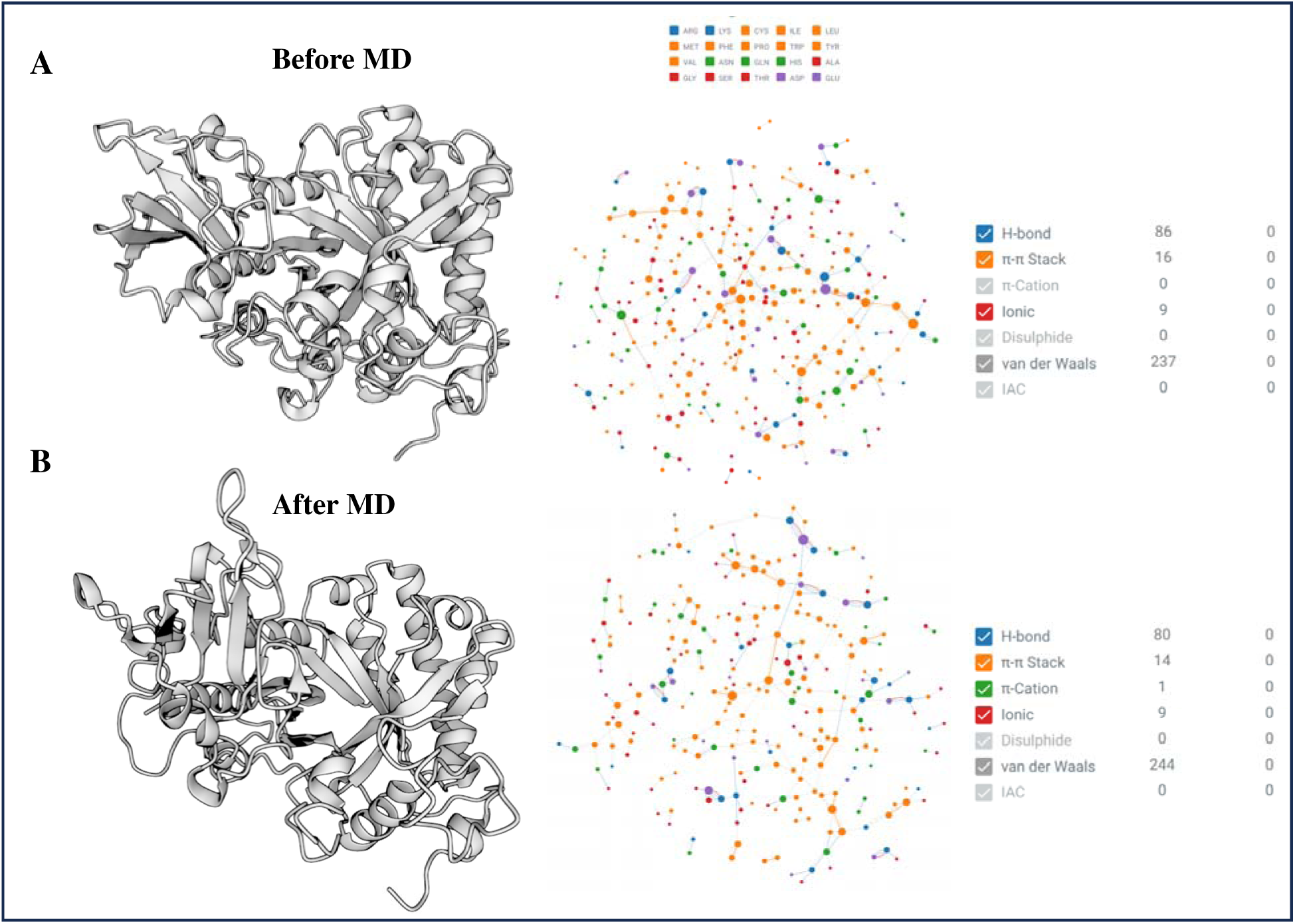
Intramolecular interaction of *PmChi*. (A) Before MD run of *PmChi* (B) Ater MD run of *PmChi*.

### 3.7 Intra-atomic interactions

Hydrophobic interactions and hydrogen bonds in the *PmChi* were the mainly structure stability providers. Total 702 (before MD) and 827 (after MD) hydrophobic bonds were present in the structure. *PmChi* had more non-polar amino acid residues in the surface of the protein Figure 6A & 6B but in the active site region had more aspartate (Asp) and glutamate (Glu) residues for executing the reaction. The non-polar rich residues located on the surface of a protein in an aqueous environment often engage in interactions with other molecules. These interactions are crucial for various biological processes. In an aqueous medium, the hydrophobic (non-polar) regions tend to avoid water and instead associate with each other or with hydrophobic regions of other molecules. This behavior can influence the folding, stability, and function of proteins(Qiao et al., 2019; Yagi et al., 2014). Hydrogen bonds are involved not only in protein-ligand interactions but also in maintaining the conformational stability of a protein, which is essential for its optimal physical properties. These bonds contribute to the proper folding and structural integrity of proteins, influencing their functionality, solubility, and resistance to denaturation under various environmental conditions (Takano et al., 1999). *PmChi* had 16 π-π stack interactions before MD and 14 π-π stack interactions after MD which maintained the protein stability, carbohydrate recognition, catalysis, and self-assembly (Shao et al., 2022). Salt bridges play a crucial role in maintaining protein stability and solubility. These interactions involve specific amino acid pairs such as His/Asp, Lys/Asp, Lys/Glu, Arg/Asp, Arg/Glu, His/Asp, and His/Glu, which are essential for forming the salt bridges. In the *PmChi* structure, among the six identified salt bridges, Lys/Asp interactions constituted 59% of the total, Arg/Asp interactions accounted for 22%, Arg/Glu interactions comprised 9%, while both His/Asp and His/Glu interactions each represented 4.5%. All salt bridges considered were less than 4.0 Å in length.

## 4. Conclusion

This study provides significant insights into the molecular characteristics of the chitinase enzyme (*PmChi*) from the mycoparasitic fungus, *P. minitans*, a widely used biocontrol agent. Through molecular modeling and simulation techniques, we have identified critical structural elements and functional residues, particularly those signifying the catalytic roles of glutamic acid (E) and aspartic acid (D) in the chitinase activity. The stability of *PmChi* during its interaction with chitotriose was confirmed, with no significant structural perturbations observed throughout the simulations. The MM/PBSA calculations proved more effective in analyzing enzyme-carbohydrate interactions, further refining our understanding of *PmChi* functionality. These findings not only enhance our comprehension of the phylogenetic and functional diversity of fungal chitinases but also pave the way for the development of improved biocontrol strategies utilizing *P. minitans*. Future research on delineation of mechanistic aspects of chitinase actions can build on these results to explore potential applications in biocontrol biotechnology.

## Acknowledgments

We thank Dr. Puneet Kumar Singh for helping in molecular simulation study.

## Author contributions

UKJ: conceptualization, investigation, data analysis, software, visualization, writing-original draft, writing-review and editing, PS: resources, supervision, writing-review and editing, NK: supervision, resources, formal analysis, writing-review and editing.

## Disclosure statement

The authors report there are no competing interests to declare.

## Notes

### Competing Interest Statement

The authors have declared no competing interest.

